# The Synergistic Effect of Antioxidant Interaction between Luteolin and Chlorogenic Acid in *Lonicera japonica*

**DOI:** 10.1101/418319

**Authors:** Meng Hsuen Hsieh, Meng Ju Hsieh, Chi-Rei Wu, Wen-Huang Peng, Ming-Tsuen Hsieh, Chia-Chang Hsieh

## Abstract

Lonicera japonica Thunb. is a flower that is used in traditional Chinese medicine to prevent the common cold. The two primary active compounds of the flower bud are luteolin, a flavonoid, and chlorogenic acid, a phenolic acid. Both active compounds have demonstrated antioxidant activity. The interactions between chemicals in a plant heavily influences its total antioxidant activity. We attempted to investigate the antioxidant interactions between the two chemicals in the plant. This study aims to investigate if the antioxidants luteolin and chlorogenic acid have a synergistic effect to inhibit free radicals when combined. A 2,2-diphenyl-1-picrylhydrazyl (DPPH•) assay was performed. The half maximal inhibitory concentration (IC50) of luteolin and chlorogenic acid were first determined and then combined at a 1:1 ratio. The combined inhibition capacity was then compared with the sum of the individual inhibition capacities. The IC50 of luteolin is 26.304 μg·ml-1 ± 0.120 μg·ml-1 while the IC50 of chlorogenic acid is 85.529 μg·ml-1 ± 4.482 μg·ml-1. The combined solution produced a free radical percentage inhibition of 77.617% ± 5.470%, more than the percentage inhibition of the separate solutions. The experiment shows that luteolin and chlorogenic acid have a synergistic effect in inhibiting DPPH free radicals.

## Introduction

*Lonicera japonica* Thunb. is a flower that is used in traditional Chinese medicine to prevent the common cold (Yang, et al. 2009). The two main constituents of the flower bud responsible for common cold prevention as active pharmaceutical compounds are luteolin and chlorogenic acid (Yuan, et al. 2014). Previous studies indicate that the two constituents both have antioxidant effects (Kono, et al. 1997; Seelinger, et al. 2008), which contribute to pharmaceutical effects of the flower bud.

The interactions between chemicals in a plant heavily influences its total antioxidant activity (Quieros, et al. 2009). Synergism of antioxidants, which refers to the interaction through which the combined antioxidant effects are greater than the sum of the individual effects (Wang, et al. 2011), has been observed in several traditional Chinese medicinal plants and is known to affect their pharmacological effects (Jain, et al. 2011; Yang, et al. 2009). However, the type of interaction in *Lonicera japonica* Thunb. has not been investigated and reported.

Because the two main active compounds of *Lonicera japonica* are antioxidants, we attempted to investigate the antioxidant interactions between luteolin and chlorogenic acid. In finding the relationship between antioxidant concentration and free radical percentage inhibition, this study aims to investigate the following question: Do the antioxidants luteolin and chlorogenic acid have a synergistic effect to inhibit free radicals when combined?

## Materials and Methods

### Antioxidant Solutions

Luteolin is soluble in water (Peng, et al. 2010) while chlorogenic acid is soluble in ethanol (Liu, et al. 2004) The antioxidant solutions of luteolin (Extrasymthèse, Rhône, France) were prepared with double-distilled water (ddH_2_O) while the antioxidant solutions of chlorogenic acid (Sigma-Aldrich, Missouri, USA) were prepared with 95% ethanol and 5% ddH_2_O.

### DPPH radical-scavenging assay

The antioxidant effects of chlorogenic acid, luteolin, and the two constituents combined will be determined via the 2,2-diphenyl-1-picrylhydrazyl (DPPH) assay, a common antioxidant assay that measures free radical scavenging capacity through spectrophotometry (Kedare, et al. 2011).

The scavenging capacity of DPPH free radical was monitored according to the methods of Wu et al. (2007), modifying this to a 96-well microtiter spectrophotometric method. 175.0 µl of DPPH• solution and 25.0 µl of antioxidant solution are added to each well. The mixture was shaken for 20 s and then left to stand at room temperature for 30 min. The absorbance of the resulting solution was read spectrophotometrically at 517 nm. The inhibition percentage (%) of radical-scavenging capacity was calculated as

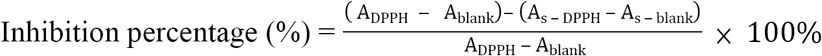

where A_DPPH_ is the absorbance of only DPPH solution, A_blank_ is the absorbance of methanol instead of DPPH solution, A_s-DPPH_ is the absorbance of DPPH solution in the presence of sample and A_s-blank_ is the absorbance of methanol in the presence of sample. The results are expressed as mmol catechin equivalents/g of sample.

Ascorbic acid, a known antioxidant, was used as a control for the DPPH assay. Ascorbic acid (Sigma-Aldrich, Missouri, USA) was used to ensure that the DPPH• solution prepared can be used to measure free radical scavenging capacity by ensuring that the resulting graph of free Radical Percentage Inhibition against Antioxidant Concentration has a linear best fit. That ascorbic acid has a consistent IC_50_ throughout the three trials is also observed to ensure that the DPPH• solution is consistent across tests.

The free radical scavenging capacity of a specific antioxidant is represented by its *half maximal inhibitory concentration* (IC_50_). IC_50_ refers to the concentration at which a specific antioxidant inhibits DPPH• in a solution by 50%. The IC_50_ is calculated from the *free radical percentage inhibition* of individual solutions using a regression curve. The concentration and volume of DPPH• solution, the volume of antioxidant solution, and the time of reaction all affect the IC_50_ of an antioxidant.

To derive a continuous graph for Free Radical Percentage Inhibition against Antioxidant Concentration, six concentrations were selected for both luteolin and chlorogenic acid. After finding the line of best fit for the graph, IC_50_, the concentration of antioxidants at which 50% of free radicals are inhibited, can be determined through the equation derived from a regression curve. Preliminary trials were performed to find the concentrations that can cover a wide range of inhibitory percentages to ensure that the regression model accurately models the data.

### Determination of Synergistic Effect

The IC_50_ of luteolin and chlorogenic acid are first separately determined using the DPPH assay. Luteolin and chlorogenic acid are then mixed at IC_50_ at a 1:1 ratio to form a combined solution. The combined solution was then diluted into 20.0, 40.0, 60.0, and 80.0% of the combined solution with both water and ethanol at a 1:1 ratio. The IC_50_ of the combined solution is then determined by graphing and expressed in % *Combined IC*_*50*_ *Solution*.

If the two antioxidants have an *additive* effect, the IC_50_ of the combined solution will remain at 100.0%, which means that the combined solution has no difference in free radical scavenging capacity compared to the separate antioxidant solutions. If the two antioxidants have an *antagonistic* effect, the calculated IC_50_ of the combined solution will be greater than 100.0%, as a higher concentration of antioxidants would be require to reduce the same number of free radicals. Conversely, if the combined solution has IC_50_ of less than 100.0%, there is a *synergistic* interaction between the two antioxidants.

Additionally, the antioxidant interaction between luteolin and chlorogenic acid can be determined with the 100.0% combined IC_50_ solution alone. If the two antioxidants have an *additive* effect, each half of the combined solution of luteolin and chlorogenic acid would inhibit 25% of free radicals in the 100.0% combined IC_50_ solution, and the resulting percentage inhibition would be exactly 50%. If the two antioxidants have a *synergistic* effect, the percentage inhibition would be more than 50%, and if the two antioxidants have an *antagonistic* effect, the percentage inhibition would be less than 50%.

### Relationship between Percentage Inhibition and Antioxidant Concentration

Per the formula of calculating Free Radical Percentage Inhibition, graphing Free Radical Percentage Inhibition (%) against Antioxidant Concentration (μg·ml^-1^) will yield a linear line of best fit. However, experimentally the free radical percentage inhibition plateaus as the antioxidant concentration inhibits close to 100% of free radicals. For this reason, the four-pair logistics function (4PL), a function that resembles a sigmoid curve, is also used by researchers to calculate the IC_50_ and different inhibitory concentrations of different substances (Sebaugh 2011). Since the center portion of the 4PL curve around the IC_50_ is linear (Sebaugh 2011), using linear regression to calculate IC_50_ may also be appropriate.

### Statistical analyses

All data are plotted using Vernier Logger Pro on a linear scale, with free radical inhibition percentage on the vertical axis and concentration on the horizontal axis. All following data sets are modeled with linear regression. The Pearson correlation coefficient (*r*), a coefficient that measures the strength of a linear relationship, and the coefficient of determination (R^2^) are both calculated for all data sets to show the suitability of using linear regression for all data sets.

*r* and R^2^ values were calculated using Vernier Logger Pro and Microsoft Excel respectively. The IC_50_ of luteolin, chlorogenic acid, and the combined solution are expressed as mean ± two standard errors of the mean (SEM).

## Results

There is a strong linear relationship between the concentration of the antioxidant solutions and the DPPH• percentage inhibition (R^2^ > 0.96) (r > 0.98), as shown in Table 3. Because there is a linear relationship between the two variables, the IC_50_ can be calculated with each linear equation obtained from each data set through linear regression.

The IC_50_ values obtained from each graph for the same antioxidant are then averaged, with the final IC_50_ expressed as mean ± two standard errors of the mean (SEM).

The calculated IC_50_ of luteolin for 25 μL of luteolin solution dissolved in water and 175.0 μL of 150.0 μmol·dm^-3^ DPPH• dissolved in ethanol is 26.304 μg·ml^-1^ ± 1.200 μg·ml^-1^, while the IC_50_ of chlorogenic acid for 25 μL of chlorogenic acid solution dissolved in ethanol and 175.0 μL of 150.0 μmol·dm^-3^ DPPH• dissolved in ethanol is 85.529 μg·ml^-1^ ± 4.482 μg·ml^-1^.

As shown in Table 2, the combined solution reaches a free radical percentage inhibition of 50% at 64.1% ± 3.2% of the combined half-maximal inhibitory concentration, which is significantly lower than 100% IC_50_, the required concentration to reach 50% free radical inhibition when the solutions are separated. This shows that a combination of luteolin and chlorogenic acid can inhibit more free radicals than the separate solutions.

**Table 3.** The Pearson correlation coefficient (*r*) and the coefficient of determination (*R*^*2*^) of each data set for the three trials performed.

| Antioxidant Solution |  | R<sup>2</sup> | *r* |
| --- | --- | --- | --- |
| Luteolin | Trial 1 | 0.9849 | 0.9937 |
|  | Trial 2 | 0.9738 | 0.9858 |
|  | Trial 3 | 0.9793 | 0.9920 |
| Chlorogenic Acid | Trial 1 | 0.9941 | 0.9971 |
|  | Trial 2 | 0.9974 | 0.9987 |
|  | Trial 3 | 0.9995 | 0.9998 |
| Combined Solution | Trial 1 | 0.9721 | 0.9859 |
|  | Trial 2 | 0.9880 | 0.9940 |
|  | Trial 3 | 0.9675 | 0.9836 |

**Table 2.** The calculated IC_50_ of the combined solution, expressed as Percentage of IC_50_ of Luteolin and Chlorogenic Acid.

| IC<sub>50</sub> of Trial 1<br>(%) | IC<sub>50</sub> of Trial 2<br>(%) | IC<sub>50</sub> of Trial 3<br>(%) | Average IC<sub>50</sub><br>(%) | Standard Deviation | Standard Error |
| --- | --- | --- | --- | --- | --- |
| 61.1 | 65.0 | 66.3 | 64.1 | 2.7 | 1.6 |

**Table 1.**
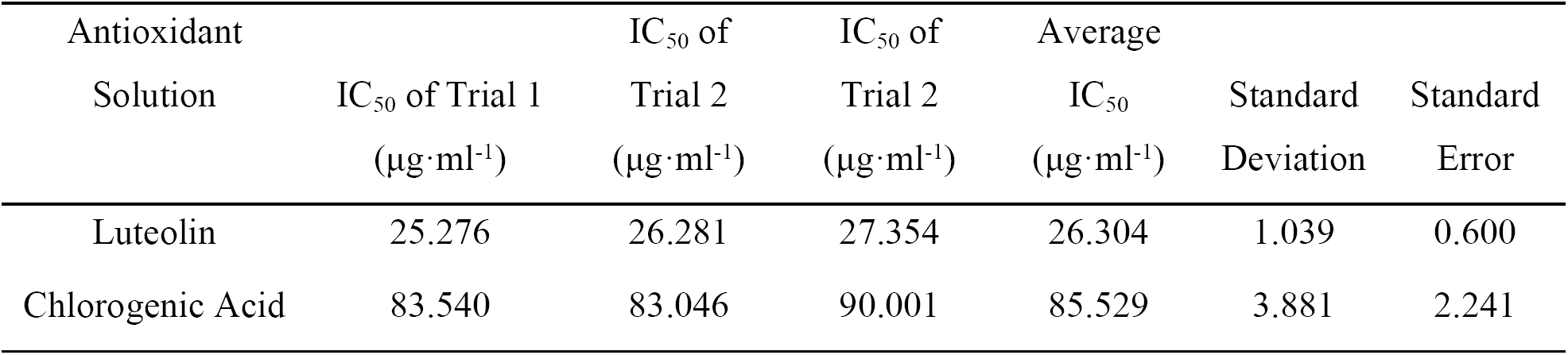
Calculated IC_50_ of Luteolin and Chlorogenic Acid from individual three trials, expressed as μg·ml^-1^

Additionally, the free radical percentage inhibition of the 100.0% combined IC_50_ solution of both luteolin and chlorogenic acid is 77.617% ± 5.470%, which is substantially higher than the percentage inhibition of the solutions at IC_50_ when separated, or 50%. That the combined solution can inhibit 77.617% ± 5.470% shows that the combined percentage inhibition exceeds the sum of the individual percentage inhibition and shows that luteolin and chlorogenic acid can inhibit more free radicals when combined.

### Types of Antioxidant Interactions

A combination of antioxidants can lead to three different types of interactions: *additive, antagonistic*, or *synergistic* (Vinson & Jang 2001). An additive effect occurs when the combined effects is equal to the sum of the individual effects, an antagonistic effect occurs when the combined of the effects is less than the sum of the individual effects, and a synergistic effect occurs when the combined effects exceed the sum of the individual effects (Vinson & Jang 2001).

## Discussion

Random error may arise from the fact that different DPPH• solutions were used for each test. DPPH• solution slowly deteriorates over time (Blois 1958). As DPPH• solution cannot be kept over long periods of time, new DPPH• solution was prepared for each test and thus the DPPH• solution used for each assay was different. Although DPPH• solution was always prepared with a concentration of 150.0 µmol·dm^-3^ for each assay performed, the concentration of each DPPH• solution created may still vary and thus have different initial absorbance for each DPPH assay performed.

Random error may also arise from the fact that the absorbance of different wells is different for the same 96-well microtiter plate. The additional absorbance of solvents is already accounted by placing “blanks” alongside wells that contained DPPH• and antioxidant solutions as illustrated in Figure 1, and through this the absorbance of each well are also simultaneously subtracted. However, error arising from the variation in the absorbance of different wells was not accounted. To account this random error, new 96-well microtiter plates were used for each DPPH assay performed to minimize the variation in absorbance.

**Figure 1.**
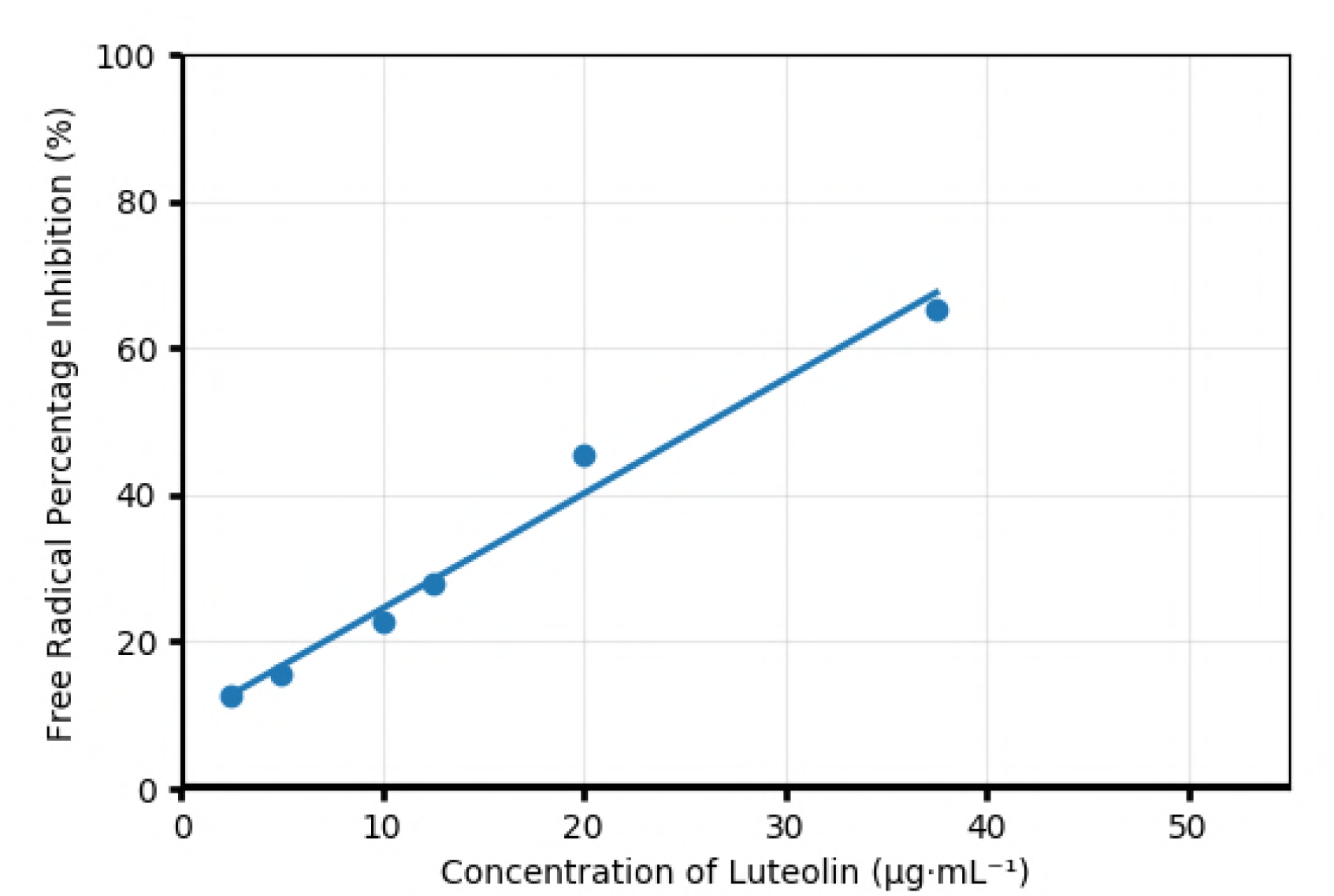
Free Radical Percentage Inhibition against Concentration of Luteolin (μg·ml^-1^)

Systematic error may arise from the fact that the absorbance of DPPH• solution increases over time. Kedare et al. suggested that the initial absorbance of DPPH• solution be under 1.0 (Kedare, et al. 2011) and that a spectrophotometric measurement of only DPPH• solution and antioxidant solvent be taken before performing the DPPH assay. For consistency, I have kept the initial absorbance of DPPH between 0.6 and 0.7 before the 30-minute reaction period. Although this confirmation was performed beforehand, the initial absorbance of DPPH• would still differ once the actual DPPH assay is performed.

From observation, the absorbance of DPPH• solution increases over time independently. The stock solution of DPPH• slowly deteriorates (Blois 1958), which may be caused by the fact that ethanol, the solvent of DPPH• used in this investigation, is volatile. 95% ethanol was used to dissolve DPPH•, and some of the solvent may evaporate during the time in which the 96-well microtiter plate is left alone for 30 minutes for the antioxidant to reduce DPPH•. The fact that DPPH• is a stable free radical (Blois 1958) and does not react with oxygen (Ionita 2005) supports this hypothesis. The evaporation of ethanol may explain the fact that seven out of the nine graphs have a positive intercept, which shows that the absorbance of the DPPH• solution may have increased because the ethanol in the DPPH• solution has evaporated over time.

Through the DPPH assay, DPPH• is reduced through hydrogen atom transfer, a mechanism that is carried out by both luteolin and chlorogenic acid. However, because there are many mechanisms through which free radicals can be reduced, most commonly hydrogen atom transfer, more than one antioxidant assays can be performed to confirm findings of a single antioxidant assays. Other antioxidant assays, such as the Ferric Reducing Ability of Plasma (FRAP) assay, which solely measures antioxidant activity with single-electron transfer (Prior, et al. 2005), may not show the same findings and results with these two antioxidants.

The goal of this research is to determine the interaction that can occur when combining luteolin and chlorogenic acid. While luteolin and chlorogenic acid have antioxidative abilities, the specific interactions between the two antioxidants in creating the synergistic effect is unknown. Present studies have been focused mostly on synergistic effects but not the specific chemical interactions as the interactions can differ widely and be quite complex (Prior, et al. 2005) Specific interactions between antioxidants can be determined by performing different antioxidant assays based on different antioxidant reduction mechanisms (Wang, et al. 2011).

## Conclusion

For all the data collected, free radical percentage inhibition and the concentration of antioxidant show a strong linear relationship (R^2^ > 0.96) (r > 0.98).

Per Figure 3 and Table 2, only 64.1% ± 3.2% of IC_50_ of luteolin and chlorogenic acid combined is required to inhibit 50% of free radicals, which is significantly lower than the 100% IC_50_ solution required for each antioxidant to inhibit the same number of free radicals separately. Furthermore, the combined solution of luteolin and chlorogenic acid at their respective IC_50_ can reduce 77.617% ± 5.470%, which exceeds the sum of the individual percentage inhibitions, or 50%. The results of the experiment conducted show that luteolin and chlorogenic acid have a synergistic effect in reducing DPPH• and may demonstrate that the pharmaceutical effects of *Lonicera japonica* Thunb. may be influenced by the synergistic interaction between the two antioxidants.

**Figure 2.**
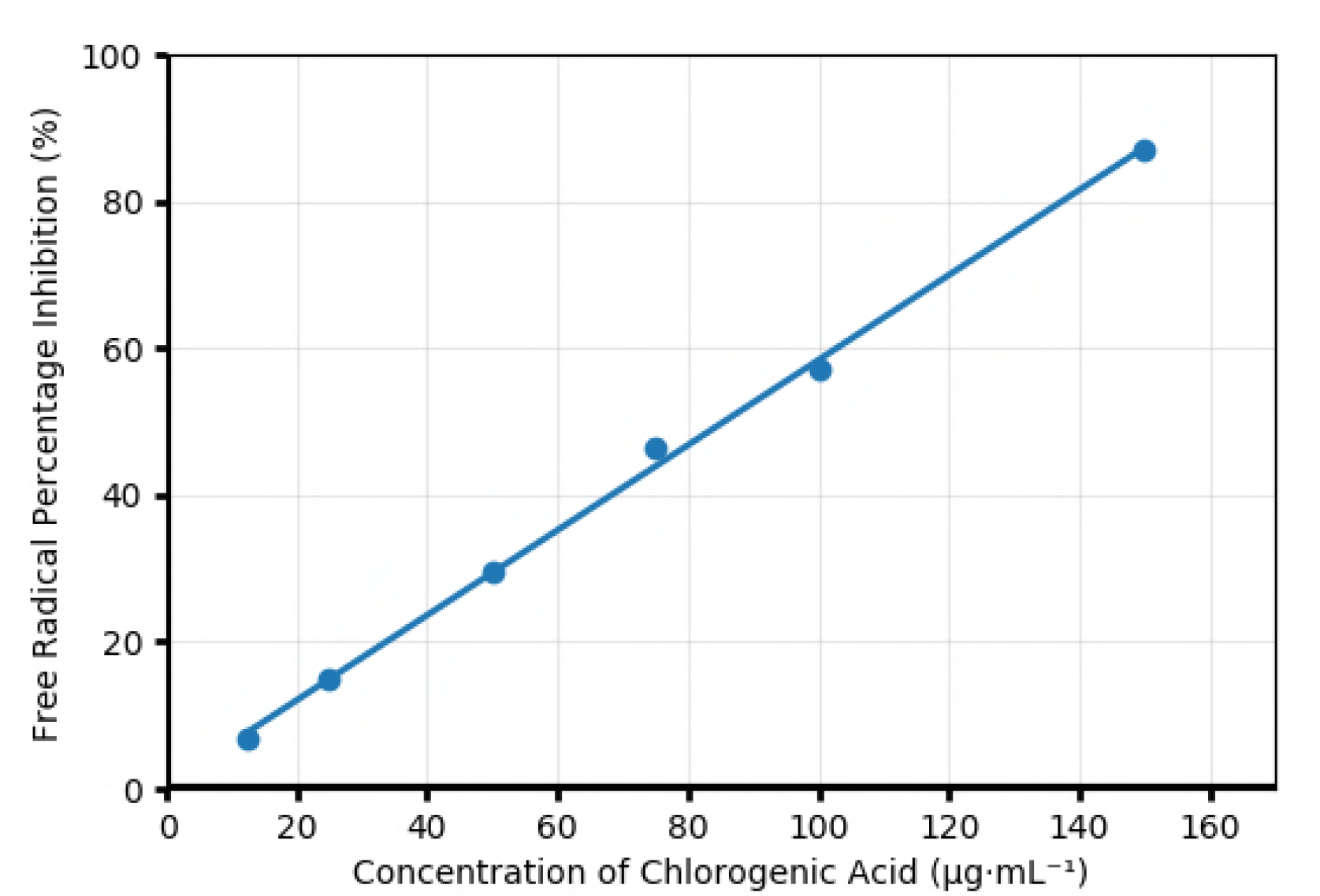
Free Radical Percentage Inhibition (%) against Concentration of Chlorogenic Acid (μg·ml^-1^)

**Figure 3.**
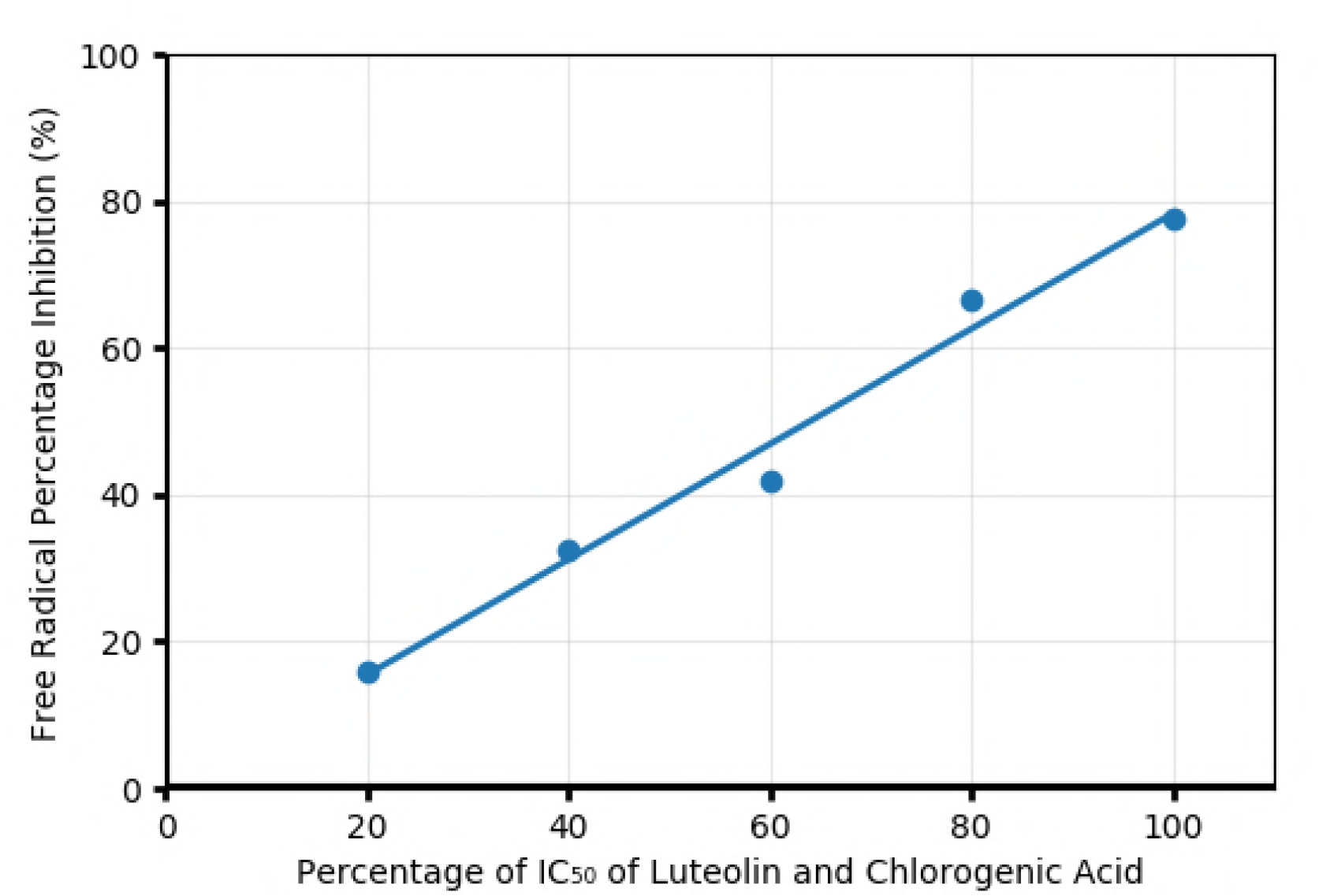
Free Radical Percentage Inhibition (%) against Percentage of IC_50_ of Luteolin and Chlorogenic Acid (%)

## References

1. Yang, Wen-Jian, Da-Peng Li, Jin-Kui Li, Ming-Hua Li, Yi-Lun Chen, and Pei-Zheng Zhang. “Synergistic Antioxidant Activities of Eight Traditional Chinese Herb Pairs.” Biol. Pharm. Bull. Biological & Pharmaceutical Bulletin 32.6 (2009): 1021–026.

2. Yuan, Yuan, Zhouyong Wang, Chao Jiang, Xumin Wang, and Luqi Huang. “Exploiting Genes and Functional Diversity of Chlorogenic Acid and Luteolin Biosyntheses in Lonicera Japonica and Their Substitutes.” Gene 534.2 (2014): 408–16.

3. Kono, Yasuhisa, Kazuo Kobayashi, Seiichi Tagawa, Koji Adachi, Akane Ueda, Yoshihiro Sawa, and Hitoshi Shibata. “Antioxidant Activity of Polyphenolics in Diets.” Biochimica Et Biophysica Acta (BBA) - General Subjects 1335.3 (1997): 335–42.

4. Seelinger, Günter, Irmgard Merfort, and Christoph Schempp. “Anti-Oxidant, Anti-Inflammatory and Anti-Allergic Activities of Luteolin.” Planta Med Planta Medica 74.14 (2008): 1667–677.

5. Queirós, Bruno, João C.m. Barreira, Ana Cristina Sarmento, and Isabel C.F.R. Ferreira. “In Search of Synergistic Effects in Antioxidant Capacity of Combined Edible Mushrooms.” International Journal of Food Sciences and Nutrition 60. Sup6 (2009): 160–72.

6. Wang, Sunan, Kelly A. Meckling, Massimo F. Marcone, Yukio Kakuda, and Rong Tsao. “Synergistic, Additive, and Antagonistic Effects of Food Mixtures on Total Antioxidant Capacities.” J. Agric. Food Chem. Journal of Agricultural and Food Chemistry 59.3 (2011): 960– 68.

7. Jain, Dheerajp, Shyams Pancholi, and Rakesh Patel. “Synergistic Antioxidant Activity of Green Tea with Some Herbs.” J Adv Pharm Tech Res Journal of Advanced Pharmaceutical Technology & Research 2.3 (2011): 177.

8. Peng, Bin, and Weidong Yan. “Solubility of Luteolin in Ethanol Water Mixed Solvents at Different Temperatures.” Journal of Chemical & Engineering Data J. Chem. Eng. Data 55.1 (2010): 583–85.

9. Liu, Junhai, and Aiyong Zuo. “Study on Technique of Extraction and Purifying Chlorogenic Acid in Eucommia Ulmoides Oliver Leaves.” Journal of Chinese Medicinal Materials 27.12 (2004): 942–46.

10. Kedare, Sagar B., and R. P. Singh. “Genesis and Development of DPPH Method of Antioxidant Assay.” J Food Sci Technol Journal of Food Science and Technology 48.4 (2011): 412–22.

11. Wu, C.-R., Huang, M.-Y., Lin, Y.-T., Ju, H.-Y., & Ching, H. (2007). Antioxidant properties of Cortex fraxini and its simple coumarins. Food Chemistry, 104, 1464–1471.

12. Sebaugh, J. L. “Guidelines for Accurate EC50/IC50 Estimation.” Pharmaceut. Statist. Pharmaceutical Statistics 10.2 (2011): 128–34.

13. Vinson, Joe A., and Jinhee Jang. “In vitro and in vivo lipoprotein antioxidant effect of a citrus extract and ascorbic acid on normal and hypercholesterolemic human subjects.” Journal of Medicinal Food 4.4 (2001): 187–192.

14. Blois, Marsden S. “Antioxidant Determinations by the Use of a Stable Free Radical.” Nature 181.4617 (1958): 1199–200.

15. Ionita, P. “Is DPPH stable free radical a good scavenger for oxygen active species.” Chem Pap 59.1 (2005): 11-16. Web.

16. Prior, Ronald L., Xianli Wu, and Karen Schaich. “Standardized Methods for the Determination of Antioxidant Capacity and Phenolics in Foods and Dietary Supplements.” J. Agric. Food Chem. Journal of Agricultural and Food Chemistry 53.10 (2005): 4290–302.

